# An Investigation of Cellulase and Epigallocatechin-3-gallate as Enhancers of Antibiotic Efficacy in Polymicrobial Biofilm Treatments

**DOI:** 10.1101/2024.10.13.617980

**Authors:** Oliver Murray

## Abstract

The treatment of bacterial infections is becoming increasingly difficult with the alarming rise of antibiotic resistance. Research has illustrated that the majority of these infections are composed of polymicrobial biofilms that can withstand antibiotic treatment to a higher degree than single-species microbial communities. Nonetheless, monospecies biofilms are still utilised for the investigation of novel combinatorial treatments, risking failure to transfer results to *in vivo* infections. This study aims to provide evidence that novel combinatorial therapies can enhance the efficacy of antibiotics in treating multispecies biofilm infections. Expanding on work within the field concerning the antibiofilm compounds epigallocatechin-3-gallate and cellulase, we sought to analyse the effects of cellulase and epigallocatechin-3-gallate, in combination with ampicillin, on polymicrobial biofilms. We pursued this by performing crystal violet assays and measuring colony-forming unit counts on environmental samples following combinatorial treatments. Statistical analysis demonstrated that cellulase significantly reduced both biofilm biomass and viable bacteria while epigallocatechin-3-gallate did not affect either quantifier. Whilst these results were not completely in line with expectations, the pronounced effect of cellulase and the epigallocatechin-3-gallate ethanol solvent were considered significant contributions to polymicrobial combinatorial treatment research. With further work in this space identifying additional therapies and examining compounds *in vivo*, there is the potential for millions of lives to be saved and the threat of a pandemic of antimicrobial resistance to be countered.

## II. Introduction

### Background

Bacteria are ubiquitous microorganisms that can be found throughout the human body, from the gastrointestinal tract to the surface of the eye [1, 2]. However, despite this range of niches, bacteria share many characteristics among an estimated 10,000 different species inhabiting the human ecosystem [3]. A universal trait of these organisms is the ability to form biofilms [4].

Biofilms are multicellular bacterial communities that display morphologically distinct attributes compared to isolated planktonic cells **(Figure 1)**. One such attribute is the production and secretion of extracellular polymeric substances (EPS), a collection of polysaccharides, proteins, lipids, and nucleic acids, into the matrix of the biofilm. This substance holds cells together in a “molecular glue”, preventing antibiotic penetration [5]. Additionally, bacteria within biofilms display altered gene expression enabling aggregation, adhesion, enhanced communication, and mechanisms to tolerate and resist antibiotics [6, 7]. These factors ultimately contribute to biofilms possessing 10-1000 times more resistance to antibiotics than their planktonic counterparts [8]. The U.S. National Institute of Health (NIH) estimates that approximately 65% of bacterial infections, and 80% of chronic bacterial infections, possess biofilms to varying extents [9]. These infections account for massive death tolls, with 1.4 million people killed by *Mycobacterium tuberculosis* in 2019 alone [10]. Considering this, and the rise of antibiotic-resistant “superbugs’’, the need to investigate biofilms and their management is of utmost importance in the face of the next global pandemic, antibiotic resistance [11].

**Figure 1.**
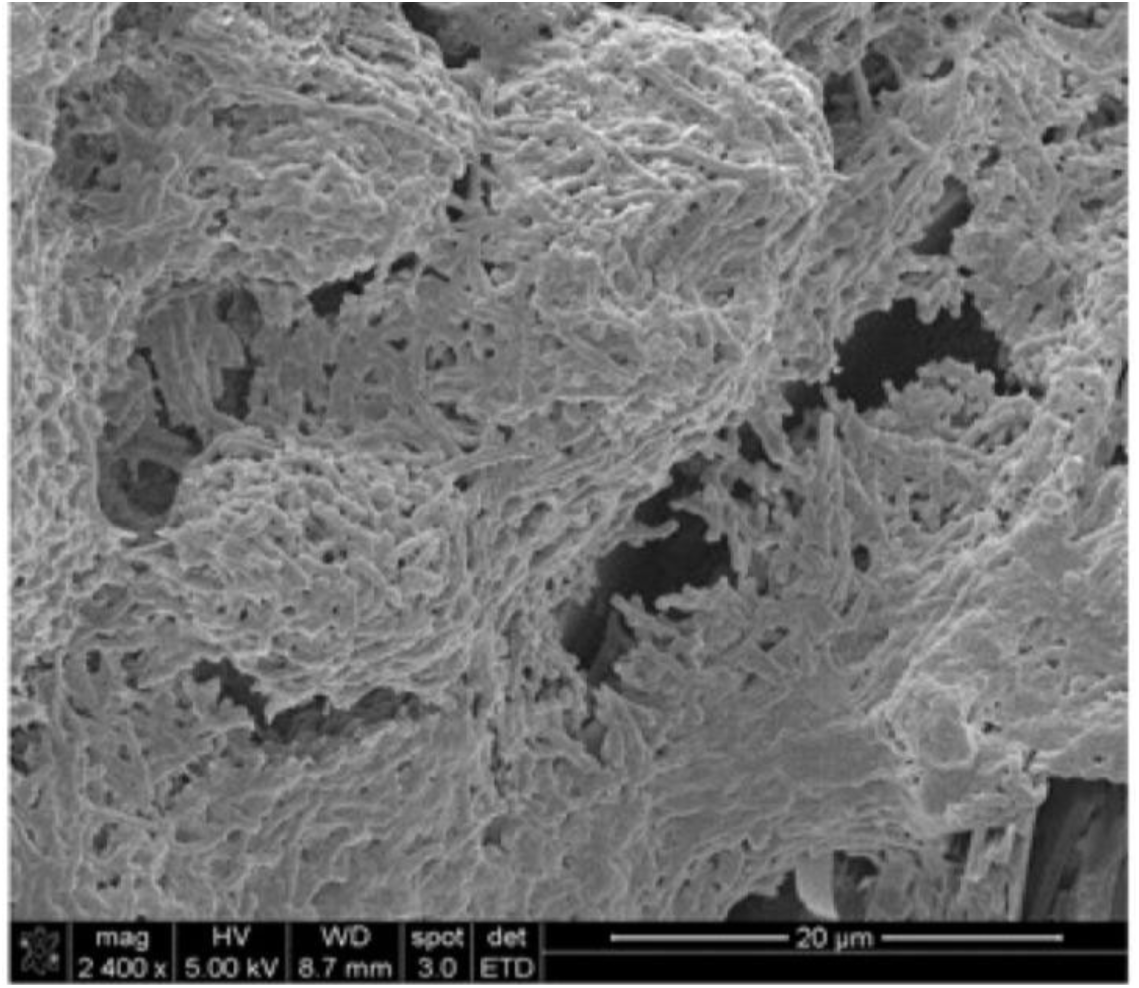
Scanning electron micrograph of *Salmonella Typh*i biofilm growing on the surface of gallbladder stones at a magnification of 2400x. Individual bacteria are closely associated and embedded within the EPS composing the extracellular matrix [12].

Biofilm investigations, both past and present, have largely focused on single-species microbial communities [13]. While these studies have provided valuable information on the key features of these microbes, their application to *in vivo* infections is hindered by their failure to examine multispecies biofilms. Polymicrobial biofilms are biofilms that contain multiple different species fulfilling different roles and niches within the microbial community. Previous work highlights that most natural biofilms, including those found within bacterial infections and chronic wounds, contain multiple different species [14]. However, clinically relevant studies still widely implement monospecies biofilms for investigation. There is evidence that bacteria display altered virulence and antimicrobial properties when grown in the presence of other species, greatly restricting the transferability of data from monospecies biofilms to those in the medical settings [15]. This is pertinent when considering that polymicrobial biofilms have a higher degree of antibiotic resistance, likely contributing to the increased mortality rates observed in multispecies infections [7, 16, 17]. Due to the increased lethality and resistance displayed by infectious polymicrobial biofilms, novel treatments must be investigated to reduce the spread and virulence of these pathogenic bodies.

Combinatorial treatments are a form of therapy that has begun to gain traction in recent decades from their usage in AIDS, malaria, and cancer treatments [18, 19, 20]. This therapy involves the delivery of multiple drugs simultaneously, targeting varying components of a pathogen, consequently reducing disease burden, and treating an infection. Due to the employment of these drugs in a concurrent fashion, the likelihood of the pathogen developing resistance to the treatment is diminished, making combinatorial treatments an attractive method to combat polymicrobial biofilm infections [21]. Indeed, work has been conducted in the literature to determine that resistance development is less common when combinatorial treatments are utilised for bacterial infections [22]. However, much of this work focuses on antibiotic combinations [23, 24]. To limit and restrict antibiotic usage, therefore preserving antibiotic efficacy, additional antimicrobial molecules should be employed in these treatments [25].

Numerous studies have investigated and proposed novel anti-biofilm agents [26]. While many of these compounds, such as antimicrobial peptides, may function well as anti-biofilm molecules, they are costly to chemically synthesise and difficult to biologically yield [27]. Considering this, cheap and readily produced anti-biofilm compounds are more suitable molecules for inclusion in combinatorial treatments. The compounds cellulase and epigallocatechin-3-gallate (EGCG) are two such molecules that fall into this category due to their wide usage in industry and ease of production [28, 29].

### Treatments

Cellulase is an enzyme produced by bacteria, fungi, plants, and animals [30]. It primarily functions to hydrolyse β -1,4 glycosidic bonds in cellulose chains, disrupting cellulose fibrils, resulting in the breakdown of cellulose-containing cell walls **(Figure 2a)**. Cellulose is one of the most abundant polysaccharides within the biofilm extracellular matrix (ECM), providing structural integrity to the microbial community **(Figure 2b/c)** [31]. As such, this is a suitable target for degradation and subsequent disruption of biofilms for enhanced antibiotic accessibility. Work within the literature has determined that cellulase is effective as a biofilm disrupting agent, suggesting that it may enhance antibiotic capabilities [32]. However, much of this work has been conducted on single-species biofilms, not accurately representing *in vivo* conditions, emphasising that work on cellulase-induced enhanced antibiotic efficacy in polymicrobial biofilms is necessary.

**Figure 2.**
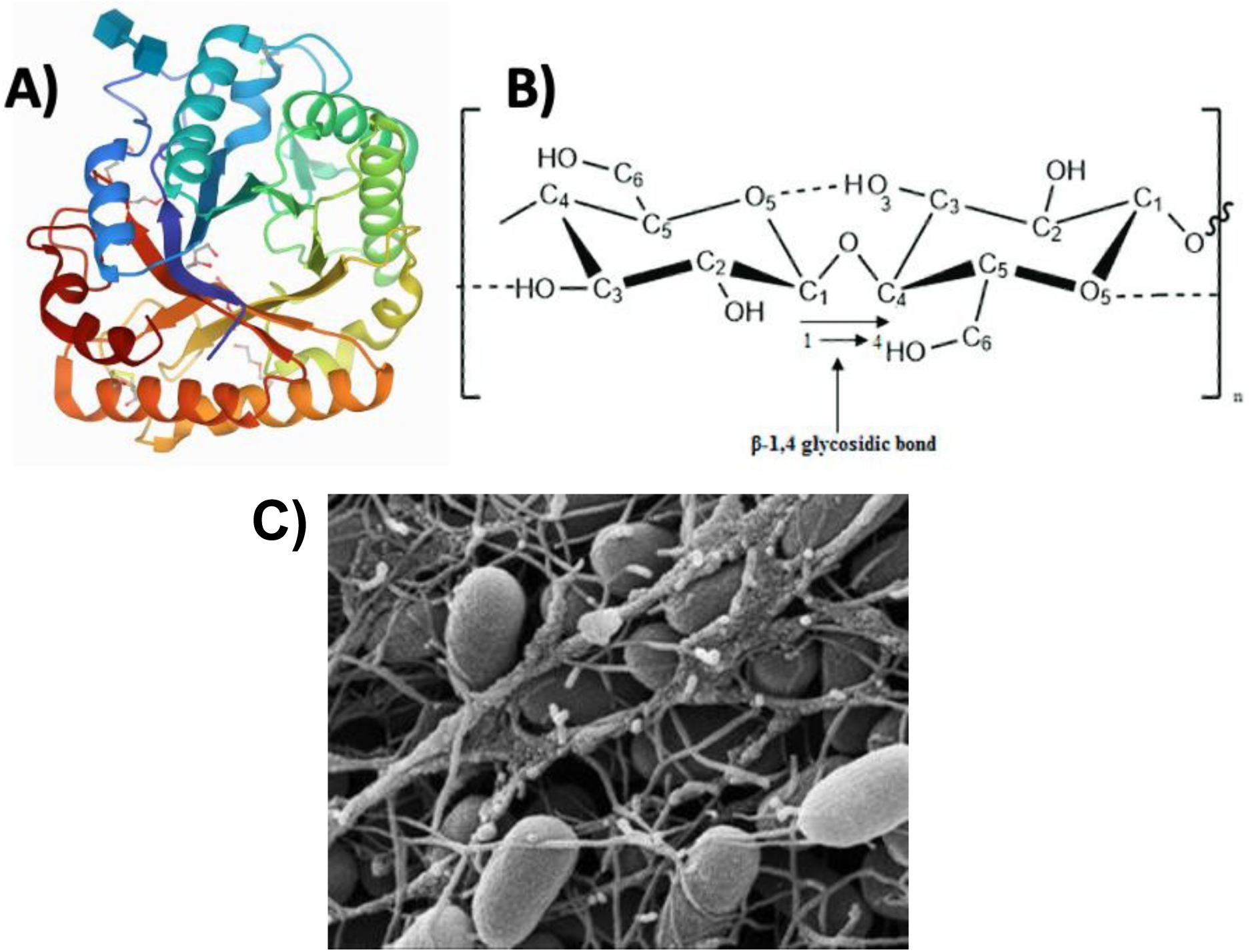
Molecular depictions of cellulase, cellulose, and *in vivo* cellulose fibrils. (A) Crystal structure of *Aspergillus niger* cellulase [33]. (B) Cellulose chain structure indicating β - 1,4 glycosidic bond for cellulase hydrolyzation [34]. (C) Scanning electron micrograph of *Escherichia coli* strain K-12 AR3110 curli-free biofilm [35]. Molecules are not to scale.

Furthermore, biofilms contain many amyloid fibrils, long repeating structural proteins composed of subunits, within the extracellular matrix [36]. One of the main amyloidogenic proteins within the ECM is the curli protein [37]. These proteins are secreted by bacteria, assemble into fibrils, and provide structural integrity to the biofilm matrix **(Figure 3A)** [38]. EGCG, a phenolic compound extracted from the tea leaves of *Camellia sinensis*, interferes with the assembly of these fibrils via two mechanisms **(Figure 3B)**. Firstly, EGCG prevents the fibrillization of curli subunits by directly interfering with their polymerisation. While this mechanism is not molecularly understood, it has been demonstrated in the lab with curli solubilisation in sodium dodecyl sulphate (SDS) [39]. Secondly, EGCG invokes the σ^E^ cell envelope stress response via the alternative RpoE sigma factor. This signal induces an sRNA, RybB, which prevents mRNA translation of a transcription factor, CsgD, involved in the transcription of curli genes [40].

**Figure 3.**
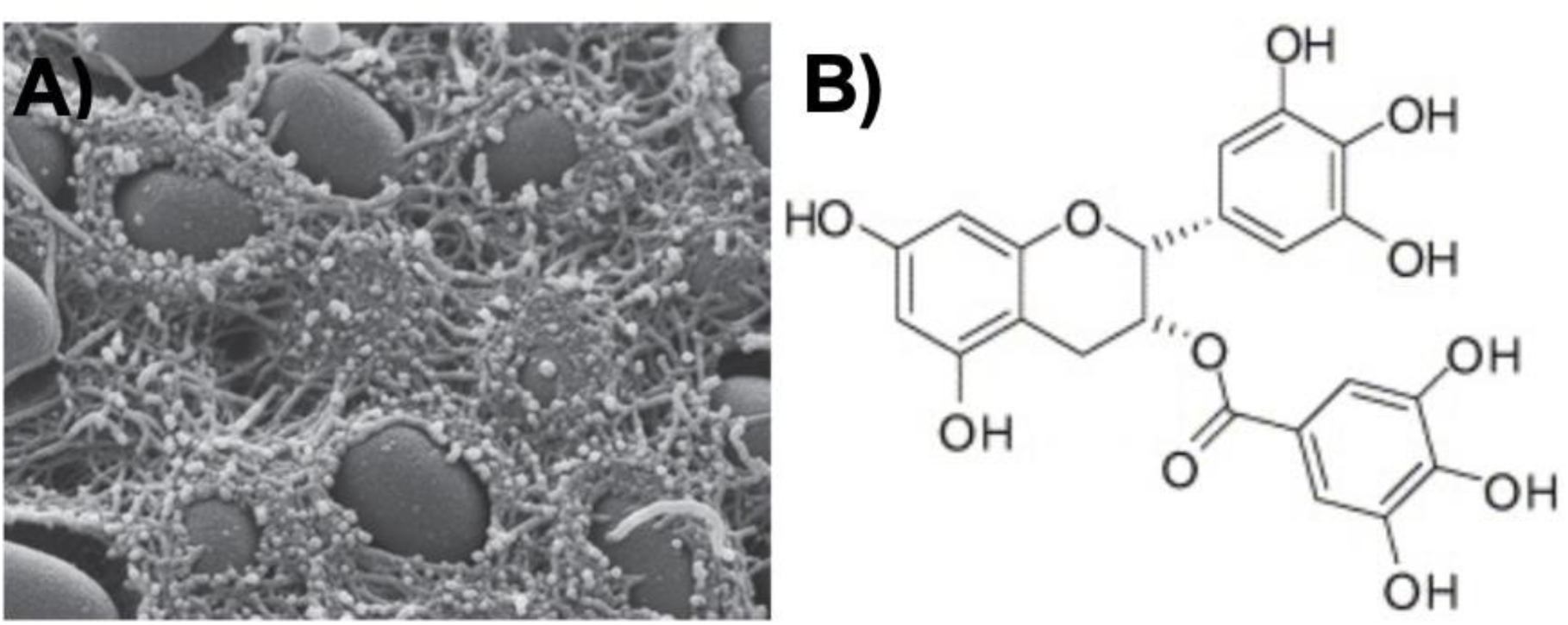
Curli biofilm amyloid fibrils and their subunit inhibitor. (A) Scanning electron micrograph of *E. coli* strain K-12 W3110 cellulose-free biofilm [37]. (B) Molecular diagram of EGCG, the curli inhibitor, depicting its reactive phenolic structure [43].

Additionally, the inhibition of CsgD prevents the transcription of a diguanylate cyclase (DgcC) involved in the activation of cellulose synthase, decreasing cellulose fibril production **(Figure 4)**. *In vitro* studies of *Streptococcus mutans* have demonstrated that EGCG can inhibit biofilm formation and production of EPS, while also reducing quorum sensing molecules and other virulence factors in *P*seudomonas *aeruginosa* via binding the LasR receptor protein [41, 42]. However, much of this work has been conducted in isolation and on single-species biofilms, leaving an undefined effect of EGCG on polymicrobial biofilms in combination with antibiotics.

**Figure 4.**
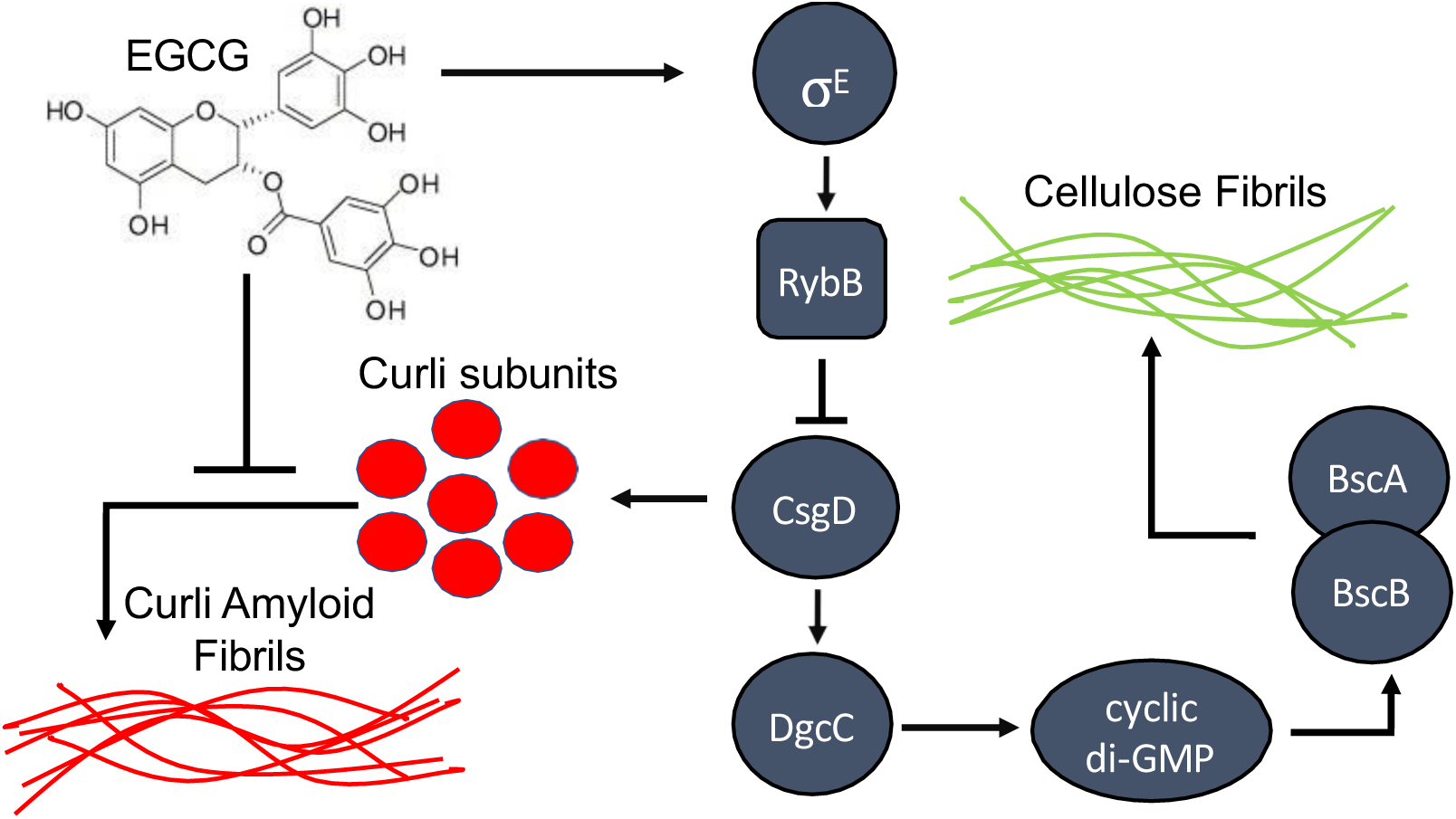
Mechanistic summary of EGCG’s inhibitory effects on the production of the biofilm ECM. Diguanylate cyclase generates c-di-GMP, a signalling molecule, which allosterically activates the cellulose synthase complex (BcsA/B). Pointed arrows indicate a mechanism of action while blunted arrows indicate a mechanism of inhibition. Molecules are not to scale.

### Experimental Question, Aim, and Methods

Performing this experiment, we aim to provide evidence that novel combinatorial therapies can enhance the efficacy of antibiotics in clearing polymicrobial biofilms, presenting a clinical solution to antibiotic-resistant bacterial infections. Examining cellulase, EGCG, and the gap within the literature concerning their clinical applicability in multispecies biofilms, it begs the question: What are the effects of cellulase and EGCG, in combination with ampicillin, on polymicrobial biofilms? We sought to answer this question by extracting an environmental multispecies biofilm to mimic a chronic wound, exposing this microbial community to our treatments, and then carrying out analytical analyses of the biofilm via crystal violet spectrophotometry assays and colony-forming unit (CFU) counts [14].

Crystal violet, also known as hexamethyl pararosaniline chloride, is a histological dye that stains negatively charged molecules [44]. The presence of peptidoglycan, a negatively charged protein component of cell walls, present in almost all bacteria, allows this dye to stain both live and dead cells. Biofilm EPS is also negatively charged from a variety of molecules containing carboxyl and hydroxyl groups [45]. Applying this dye, allowing biomass adherence, rinsing excess, and removing adhered dye, allows for quantification of biomass via absorbance measurements in a spectrophotometer. Considering this, crystal violet is a suitable measurement of the overall biomass of biofilms.

CFU counts are measurements of viable bacteria within a sample. These are quantified following treatments, biofilm disruption, extraction of supernatant, serial dilution, spread plating, and incubation. After an adequate incubation period, colonies are observed on agar plates and counted **(Figure 5)**. Colony counts are converted to CFU/mL with the following formula to allow for treatment comparison analysis of viable bacteria in biofilm samples after treatments:

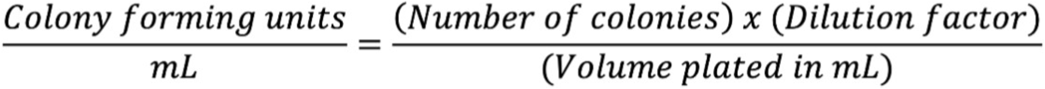

**Figure 5.**
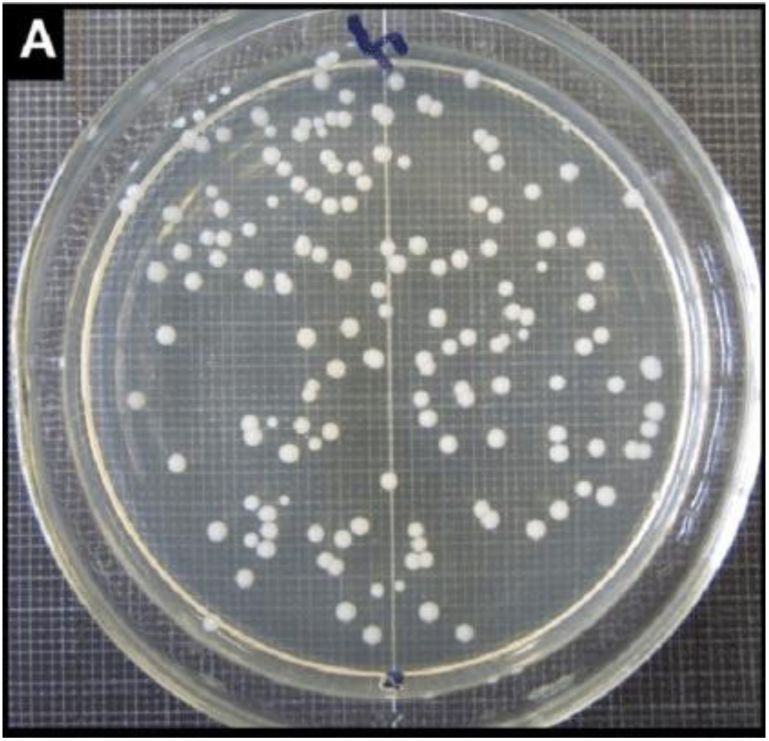
*Staphylococcus aureus* colonies after 24 hours of growth on an agar plate [46]. Individual colonies represent growth from a single bacterium.

### Hypotheses

Utilising crystal violet to quantify biomass and CFU to determine viable bacterial loads, we seek to determine the validity of two hypotheses:

*Ha*1: EGCG in combination with ampicillin will significantly reduce crystal violet absorbance and CFU, compared to the ampicillin treatment in isolation, on polymicrobial biofilms.
*Ha*2: Cellulase in combination with ampicillin will significantly reduce crystal violet absorbance and CFU, compared to the ampicillin treatment in isolation, on polymicrobial biofilms.

We then evaluate our results through comparative statistical analyses, a discussion of each of the treatments within the context of the literature, and consideration of any experimental limitations, to determine if EGCG and/or cellulase serve as an enhancer of antibiotic efficacy. Following this, we recommend further work that may be conducted to offer greater applicability to our results and expand the field of combinatorial treatments.

## IV. Materials and Methods

All LB broth and agar were produced from their powdered forms and diluted with distilled water. Phosphate-buffered saline (PBS), acetic acid, and crystal violet solution were provided courtesy of The University of Southampton Microbiology Laboratory.

### Sample Collection, Cultivation, and Selection

A water sample was collected from The Valley Gardens’ ponds on the University of Southampton Highfield Campus. The sample was taken back to the lab and a serial dilution was performed in 1/10 LB broth to remove any remaining debris, select a suitable level of growth, and induce biofilm formation via nutrient deprivation [47]. Next, 5.0 mL from each dilution was deposited into a well from a six-well plate, each containing a glass coverslip for biofilm growth.

After 3 days of incubation within the 25 °C chamber, the excess broth was dispensed, and samples were rinsed twice with PBS. Following this, the glass coverslips were examined underneath a light microscope at a magnification of x100 for selection. The 10^-2^ dilution was chosen for moderate growth.

The 10^-2^ dilution coverslip was reinserted into its well, inoculated with 3.0 mL of 1/10 LB broth, sheared with a cotton bud, and 100 μL of supernatant was distributed across 6 wells with coverslips. After an additional 3 days of growth at 25 °C, the excess broth was dispensed, and samples were rinsed twice with PBS. This procedure was then repeated to 42 wells to produce the number of biofilms required for experimentation.

### Treatment Preparation

*Aspergillus niger* powdered cellulase, purchased from Sigma Aldrich, was dissolved in acetate buffer, produced following the AAT Bioquest protocol, to yield a concentration of 10.4 mg/mL [48]. This itself was then diluted in 1/10 LB broth to 10 mg/mL such that 9.4 enzymatic units were utilised in coherence with prior work [49].

EGCG was dissolved in ethanol [50]. This solution was then dispensed into 1/10 LB broth to produce a concentration of 100 μg/mL of EGCG as a median dosage of the lipid derivative of EGCG, EGCG-stearate, within the literature [51].

Ampicillin was used as the antibiotic in this experiment. From a stock solution contained within Dr Dumont’s lab, a 50 μg/mL dosage was prepared in 1/10 LB broth. This served as an easily producible median value to the 20 μg/mL and 200 μg/mL ampicillin dosages utilised in a previous study that we conducted [52].

### Treatment Applications

5.0 mL of treatments were applied in their diluted states and given to biofilms on coverslips within well plates. Treatments for crystal violet and colony-forming unit analysis were both applied for 24 hours to achieve consistent results. Different combinations of treatments were applied for a comprehensive analysis, including a control of 1/10 LB broth. Following treatment completion, wells were emptied and rinsed twice with PBS.

### LB Agar Plate Preparation

LB agar plates were produced aseptically using flaming techniques to avoid contamination and were left to sit on the workbench overnight.

### Crystal Violet Staining and Quantification

0.1 % v/v crystal violet staining solution was provided and applied to glass coverslips for 20 minutes. Following exposure, 3.0 mL of 30% acetic acid solution was applied to each well for 20 minutes and placed onto a see-saw rocker. After 20 minutes, 1.0 mL was extracted from each well and placed into a cuvette for spectrophotometer analysis at 584 nm.

### Colony Forming Units: Plating and Evaluation

After rinsing samples with PBS twice, wells were inoculated with 5.0 mL of 1/10 LB broth and biofilms were gently sheared with a cotton bud. A serial dilution was then performed. 50 μL was aseptically spread plated from each dilution of each treatment and plates were left to grow for 3 days to achieve optimal growth. Only plates with 20-200 colonies were selected for interpretation to decrease the effect of low counts on statistical analysis and maintain a degree of accuracy in colony counting.

## V. Results

### Crystal Violet Spectrophotometry

To develop an understanding of combinatorial treatment effects, we performed a crystal violet stain on biofilms after 24 hours of exposure to their respective treatments. Following this procedure, the rinsing of excess dye, and the addition of acetic acid to the wells, we analysed the supernatant in a spectrophotometer **(Figure 6)**. Applying this method allowed for the comparison of remaining biomass in each sample following treatment exposure. We examined a multitude of treatment combinations to determine if any synergistic or antagonistic effects were present.

**Figure 6.**
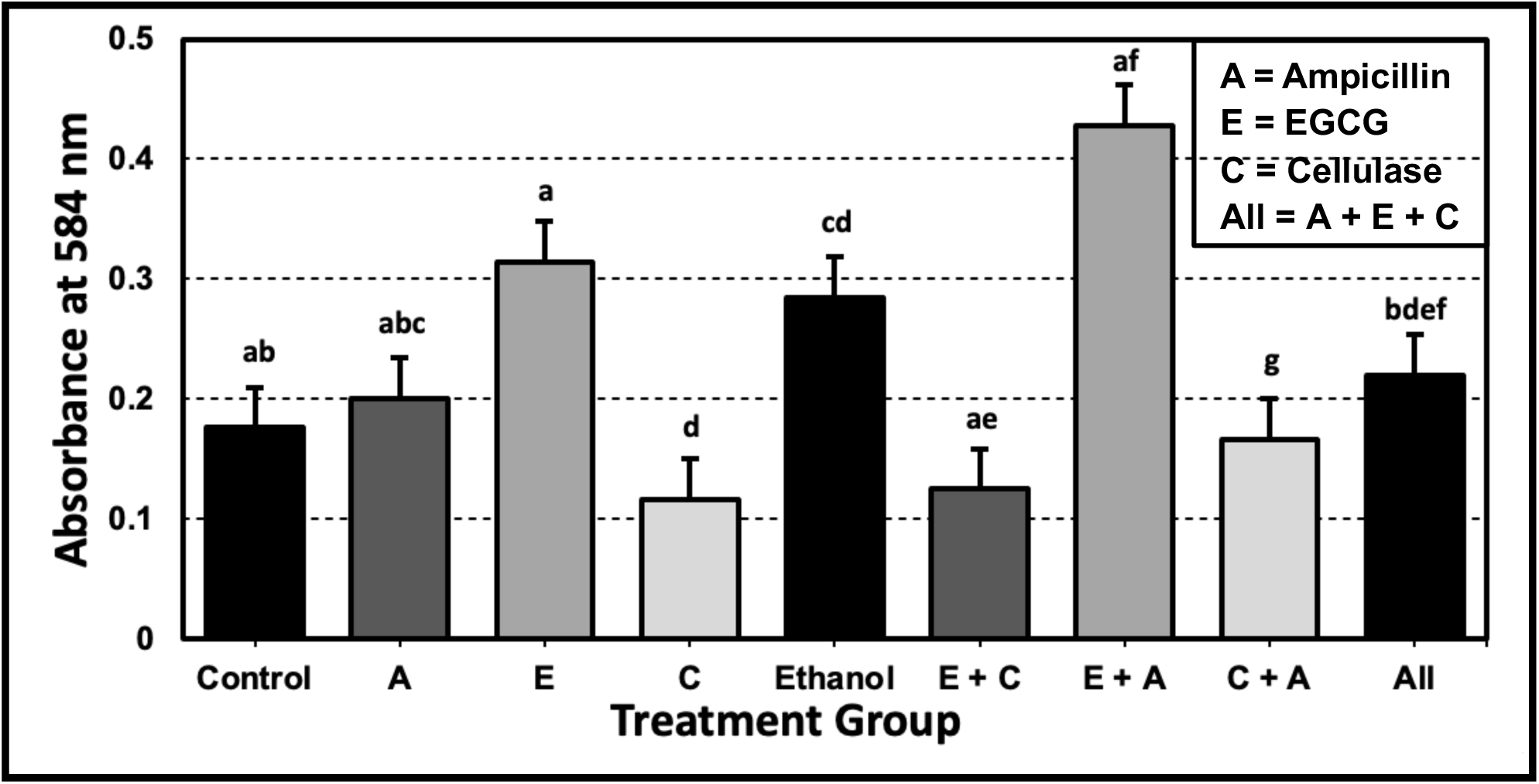
Spectrophotometer absorbance of crystal violet solution displaced from biofilm samples following treatments. Ethanol was examined as a positive control for the solvent of EGCG. Error bars indicate SEM. Differing lowercase letters indicate significantly different absorbance values determined by Tukey’s Test (*p* < 0.05). For all spectrophotometry data, please view the Appendix **(Table A1)**.

Visibly the data appeared to contain a large degree of variation between treatment groups. To further analyse these results, we performed a one-way analysis of variance (ANOVA) to investigate if there was any statistical difference in absorbance values between treatments.

Implementing this, we determined that overall, the treatment groups had significantly different absorbance values (*F*_8,18_ = 27.349, *p* < 0.0001). However, this analysis failed to determine which specific treatment groups were statistically different. To address this issue, we performed Tukey’s Test. This test revealed many statistically significant differences between treatments, the most notable of which we encapsulated in table format **(Table 1)**. Further, this test highlighted that there was no difference between the control, ampicillin, EGCG, or the EGCG and ampicillin combination treatment.

**Table 1.** Admissible statistically different absorbance results from Tukey’s Test, indicating biomass differences. Additional results were excluded from the table due to lack of study applicability. For all of Tukey’s Test results, please view the Appendix **(Table A2 and A3)**.

| Treatments | Q statistic | p-value |
| --- | --- | --- |
| Control vs Ethanol | 5.645 | $p < 0.05$ |
| Control vs Cellulase and Ampicillin | 13.090 | $p < 0.01$ |
| Ampicillin vs Cellulase | 5.905 | $p < 0.05$ |
| Ampicillin vs Cellulase and Ampicillin | 11.826 | $p < 0.01$ |
| EGCG vs Cellulase | 10.268 | $p < 0.01$ |
| EGCG vs Ethanol | 8.744 | $p < 0.01$ |
| Cellulase vs EGCG and Cellulase | 9.835 | $p < 0.01$ |
| Cellulase vs Cellulase and Ampicillin | 5.922 | $p < 0.05$ |
| Cellulase and Ampicillin vs All | 10.822 | $p < 0.01$ |

Considering that higher absorbance equates to higher biomass in samples, these results suggest five findings. Firstly, cellulase reduced biomass compared to the ampicillin treatment by 41.86%. Secondly, the ampicillin and cellulase combinatorial treatment, while not effective as cellulase alone, still reduced biofilm growth by 20.64% compared to the ampicillin treatment. Thirdly, EGCG increased biofilm growth by 32.26% in combination with cellulase and ampicillin compared to cellulase and ampicillin alone. Additionally, the ethanol-treated biofilm biomass was 61.62% greater than that of the control, while EGCG treated biofilms were 10.29% greater than ethanol. Finally, ampicillin, EGCG, and the combinatorial treatment of ampicillin and EGCG had no significant effect on the biofilm biomass compared to the control treatment.

### Colony Forming Units

While the crystal violet data was insightful into the effect of treatments on biomass, it did not distinguish between live and dead bacteria. To accomplish this distinction, we performed a CFU analysis to gain further insight into the outcome of the treatments on surviving bacterial counts within the biofilms **(Figure 7)**. The calculation was simplified with all usable plate counts having a dilution factor of 10^-5^. In terms of treatments, a reduced selection was implemented compared to the crystal violet analysis to concentrate the investigation on the effect of EGCG and cellulase on the efficacy of ampicillin treatment.

**Figure 7.**
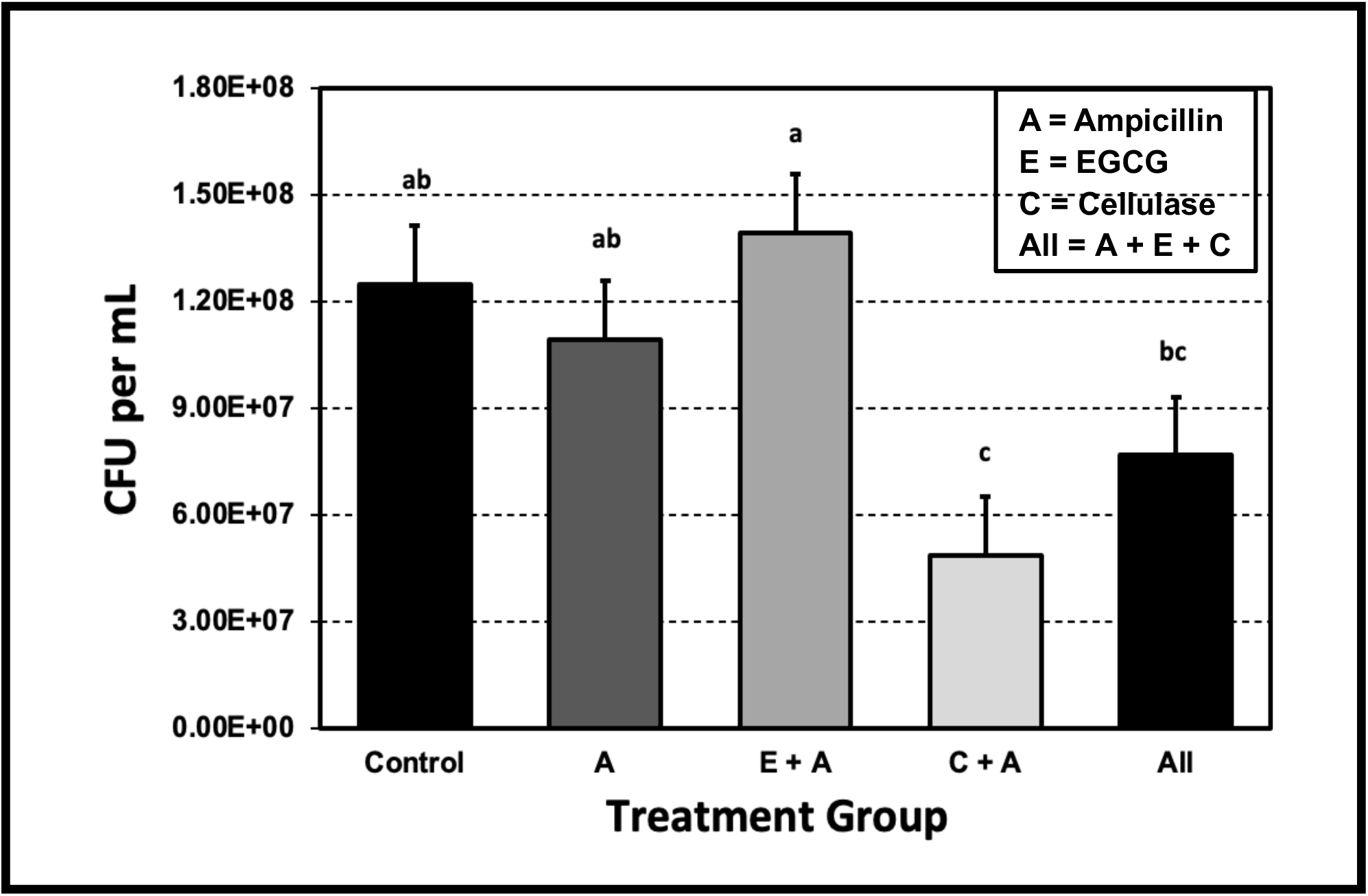
Colony-forming unit (CFU) counts per mL of biofilms exposed to treatment groups. Error bars indicate SEM. Differing lowercase letters indicate significantly different CFU/mL determined by Tukey’s Test (*p* < 0.05). For all CFU data, please view the Appendix **(Table A4, A5, and A6)**.

First impressions of data appeared to indicate that the cellulase and ampicillin combinatorial treatment contained fewer CFU/mL than the ampicillin treatment. To confirm this observation, a One-Way ANOVA was performed, demonstrating there was a significant difference between treatment groups (*F*_4,10_ = 9.002, *p* < 0.005). We followed this analysis with Tukey’s Test to determine specific significantly different treatment groups. This investigation revealed there was a significant difference between ampicillin and the combinatorial cellulase and ampicillin treatment, with the combinatorial treatment containing 53% fewer CFU/mL. Further, there was no significant change in CFU/mL between the control, the ampicillin treatment, or the combinatorial EGCG and ampicillin treatment. The treatment containing ampicillin, EGCG, and cellulase was found to not statistically differ from the control, ampicillin, or combinatorial cellulase and ampicillin treatment.

These results indicate that cellulase had the effect of decreasing colony-forming units when combined with ampicillin as opposed to the ampicillin treatment in isolation. Further, the statistical data proposes that neither ampicillin nor the combinatorial EGCG and ampicillin treatment had an effect on the colony-forming units present within the biofilms.

## VI. Discussion

Throughout our experiment, it was evident that ampicillin was ineffective at reducing biofilm biomass and CFU. Utilising statistical analyses, we found that there was no statistically significant difference between the control and the ampicillin treatment. Such evidence contributes to a growing field, emphasising that multispecies biofilms deserve greater attention and experimental investigation within the scientific community due to their antibiotic resistance qualities. Building on this concept, here we will discuss our results for EGCG and cellulase in the context of the literature, the limitations of our study, and recommendations for further work to continue to explore the field of combinatorial treatments.

### Epigallocatechin-3-gallate

We hypothesised that EGCG, in combination with the ampicillin antibiotic, would decrease polymicrobial biofilm biomass and CFU based on previous work within the literature. For example, studies into the effects of EGCG on *E. coli* found that the inhibition of curli fibres and decrease of the cellulose production control gene, *DgcC*, by EGCG resulted in the elimination of the biofilm matrix [40]. Additionally, *in vivo* polymicrobial cystic fibrosis mouse models determined that nebulized EGCG reduced CFU/mL significantly, suggesting a bactericidal effect [53]. This concept was enforced by the synergistic effects of EGCG and carbapenems (imipenem, panipenem, and meropenem) on methicillin-resistant *S. aureus* (MRSA) inhibiting growth [54].

However, in our crystal violet assay, we found that EGCG and the combinatorial EGCG and ampicillin treatment did not have any effect on biofilm growth/biomass when contrasted with the control. Further, when compared to both the cellulase treatment and cellulase ampicillin combination therapy, the addition of EGCG to the treatments significantly increased the biomass. These results indicate that EGCG is not only ineffective at reducing multispecies biofilm growth but that in certain instances it may counteract treatments and promote biofilm proliferation. Our CFU analysis confirmed this result in that the combinatorial EGCG and ampicillin treatment was ineffective compared to ampicillin and the control in reducing viable bacteria within the polymicrobial biofilms.

The literature highlights that EGCG is species-specific in its inhibition of curli amyloid fibre formation, and induction of the σ^E^ cell envelope stress response [55, 56]. Further, many previous studies demonstrating the effectiveness of EGCG in biofilm treatments are examinations of monospecies biofilms such as *E. coli* [40]. Our research contributes directly to this information, providing evidence that EGCG possesses a lack of efficacy when interacting with nonspecific multispecies biofilm communities. While this is clear evidence that EGCG is not effective as a broad range antibiofilm agent, there is additional work that suggests EGCG may function as a nutritional carbon source in certain species, offering a further explanation for its lack of overall effect due to its metabolism [57]. As such, we gather that our results demonstrating minimal or positive effects of EGCG exposure are likely due to a range of factors involving the complexities of multispecies biofilms, different bacterial species to those observed to respond to EGCG, and differing ECM constituents [58].

Such evidence has implications for further work and contributes to a growing consensus of research that argues EGCG is not a cure-all solution to biofilms. Indeed, the literature is unsure about the treatment capacity that EGCG possesses, with many early studies supporting its usage to counter the rise of antibiotic resistance [54, 59]. However, recent work suggests that the ability of EGCG to interfere with biofilms is more complex, with some studies describing how EGCG may enhance vancomycin, oxacillin, and ampicillin antibiotic tolerance in a similar manner to our results, therefore promoting biofilm growth [60, 61]. Interestingly, an additional factor that may have contributed to EGCG promoting biofilm growth within our study was the EGCG ethanol solvent, which we determined promoted biofilm growth. The literature supports this postulation with studies determining that ethanol has a positive effect on biofilm monoculture growth of *S. aureus* and *epidermidis* [62]. Studies have demonstrated that ethanol exposure downregulates the Spx transcription factor, decreasing the transcription of the intercellular adhesion (*ica*) operon repressor (*icaR*). This results in *ica* promoting intercellular adhesion polysaccharide production, increasing biofilm adherence and growth [63]. While ethanol did increase biofilm biomass, the EGCG treatment had 10.29% more biomass than the ethanol treatment, suggesting that while ethanol does contribute to the unproductive and promotional effects of EGCG it does not account entirely for this result.

Considering that our study is the first published experiment examining EGCG on environmental biofilms to replicate polymicrobial infections, the possible lack of clinically relevant species, including those that have been identified to be affected by EGCG, may account for our data. Broadly speaking, pond water microbiota consists of a diverse set of species such as *E. coli*, *Enterobacter aerogenes*, *Klebsiella pneumoniae*, *Pseudomonas spp*., *Bacillus spp.*, *Listeria monocytogenes*, and *Staphylococcus spp.* [64]. Due to laboratory constraints, we were unable to determine which species we had present within our samples. As such, our data may not be significantly transferable to polymicrobial biofilm treatment as different biofilms may contain differing compositions of organisms and ECM constituents. Although, we argue that despite composing an unknown microbiota, there is evidence that many of the bacteria involved in chronic wounds, *P. aeruginosa*, *S. aureus*, *K. pneumoniae*, *Enterobacter spp.*, and coagulase-negative *Staphylococci spp.*, are commonly found in pond water [64, 65]. While the species within our polymicrobial biofilms are unknown, we suggest clinically relevant bacteria may be present within the sample, offering validation to our results. Additionally, as discussed, the vast majority of studies into the effectiveness of EGCG have been performed on single-species biofilms. Therefore, we infer that our usage of a multispecies biofilm contributed to our observations that EGCG was ineffective and promoted the growth of the multispecies biofilm biomass. With this in mind, we conclude that EGCG does not enhance ampicillin efficacy in multispecies biofilm treatments.

### Cellulase

Previous studies indicate that cellulase in isolation cannot inhibit biofilm formation but may do so in combination with antibiotics [49]. Indeed, further work conducted determined that the interaction of cellulase and ceftazidime induced enhanced single-species biofilm disruption of the chronic wound pathogen *P. aeruginosa* [66]. These studies served as the basis of our hypothesis that cellulase, in combination with ampicillin, would decrease polymicrobial biofilm biomass and CFU.

Examining our spectrophotometry data revealed that while our combinatorial treatment of cellulase and ampicillin was not significantly as effective as cellulase alone, it was able to decrease the absorbance, and thus biomass of the biofilms compared with the ampicillin treatment. This result was reinforced by our CFU data which indicated that cellulase decreased viable bacterial counts when combined with ampicillin as opposed to the ampicillin treatment in isolation. These methods implicate that cellulase in combination with ampicillin may reduce both biomass and viable bacteria compared to the standard ampicillin treatment. Work within the literature on a range of different monospecies biofilms supports our findings, providing evidence that EPS cellulose is a common component of bacterial biofilms, despite species possessing vastly differing EPS constituents [67]. This extensive cellulose incidence explains why cellulase can polymicrobial communities. Further, recent studies conducted with mouse models have determined that cellulase may possess enhanced clearance capabilities against *P. aeruginosa* established polymicrobial biofilms, lending additional support to our investigation [15].

In contrast to the literature, our cellulase treatment was able to independently reduce biofilm biomass, thus demonstrating the potential to prevent biofilm formation. For example, an examination of the cystic fibrosis pathogen *Burkholderia cepacia* determined that while cellulase did possess a general antimicrobial property, it was not able to affect growth, and thus biomass [68]. Our observation that cellulase can reduce bacterial growth in isolation against polymicrobial biofilms directly contradicts this, making it the first discovery of its kind. A possible explanation for this occurrence is our procedural removal of the supernatant from the wells.

Cellulase functions by cleaving cellulose chains, disrupting the cellulose fibrils that bind the ECM together, resulting in fragmentation of the biofilm structure [69]. Such fragmentation may have caused biomass and viable bacteria to be released into the supernatant of the wells, not being recorded within the data, and reducing observed values. While this does affect our isolated cellulase treatment, the impact in combination with ampicillin is minimal due to the nature of both cellulase and ampicillin acting upon the exposed layer of the biofilm due to poor antimicrobial penetration [70]. This results in the removed bacteria having a greater likelihood of already being incapacitated by ampicillin. Additionally, it must be considered that viable bacteria removed from the ECM are easier for ampicillin, or any other antimicrobials, to interact with due to the lack of ECM restrictions. While this limitation must be considered for absorbance values relating to biomass, it has a diminished effect on CFU. Considering this, further work needs to be performed to investigate isolated cellulase antimicrobial properties on polymicrobial biofilms.

Our spectrophotometry result indicated that cellulase and ampicillin were not as effective as cellulase in isolation in the reduction of biofilm biomass. There is evidence that antibiotic exposure may induce biofilm formation with subinhibitory concentrations [71]. This mechanism, whilst not fully understood, is believed to occur via increased expression of the *cidA* gene involved in regulating cell death, resulting in increased biofilm thickness, and preventing antibiotic penetration [72]. Further studies determined that antibiotics can also trigger cell envelope stress responses to induce biofilm formation in *S. mutans* via regulation of the *atlA* and *rgp* genes modulating cell wall components [73]. However, examination of treatment recommendations across a range of medicinal practices, including the CDC, suggests that confirmation of antibiotic-resistant infections is treated with second and third antibiotic series, particularly ‘last-resort’ carbapenem class antibiotics [74, 75]. Essentially, despite the presence of antibiotic resistance, antibiotics will be utilised in one format or another, validating the clinical usage of our combinatorial treatment. It must also be considered that bacterial infections and chronic wounds have fewer nutrients, a vast quantity of inflammatory molecules, extracellular proteases, and increased host cellular contents [76]. This contrasts with the relatively nutrient-rich, noninflammatory, protease-free, *in vitro* assay conditions we employed. Furthermore, our bacteria do not need to contend with an immune system attempting to clear the polymicrobial infections *in vivo*. While our experimental setup does not mirror what occurs *in vivo*, it demonstrates that even in optimal growing conditions our cellulase and ampicillin combinatorial treatment is extremely effective. Therefore, this treatment likely possesses greater application to *in vivo* therapies by working with the immune system to clear the infection. Considering this, our spectrophotometry and CFU results, as well as an in-depth analysis of the literature, we conclude that cellulase enhances ampicillin efficacy in multispecies biofilm treatments.

### Recommendations for Further Work

Our study has provided evidence that cellulase can be utilised to increase the efficacy of ampicillin in clearing polymicrobial biofilms, while EGCG struggles to do so. However, more research must be conducted to validate our results in light of the intended application of our work within the clinic.

Firstly, we are unsure of the bacterial species present within our biofilm. As we have described, there is potential for similar species to be present in our pond water sample and chronic wound polymicrobial biofilms. However, we advise that identification via biochemical tests, antibody targeting, and genetic analyses is necessary to increase the applicability of our data [77].

Secondly, additional confirmation of the enhanced efficacy of our cellulase and ampicillin treatment is necessary. This should be carried out via a live/dead stain with SYTO9 and propidium iodide to analyse the proportion of life to dead cells within the biofilms [78].

Additionally, we would recommend taking this a step further with an analysis of the supernatant that we withdrew to address our procedural experimental limitation of isolated cellulase efficacy. Employing these means, it can be concluded if our treatment is resulting in biofilm fragmentation followed by cell death, or simply biofilm displacement into the supernatant.

Thirdly, experimentation on ethically extracted polymicrobial biofilms from patients with chronic wounds would further facilitate the introduction of our treatment into the clinic by validating our data with known pathogens. If validated, we believe that our cellulase and ampicillin treatment would be most beneficial in clearing superficial chronic infections due to ease of application in solution, as well as the concentration of its effect. In particular, the addition of cellulase to antibiotic ointments such as mupirocin would be paramount to combatting its exceedingly high rate of resistance development [79].

Further, the employment of a different solvent to ethanol is necessary to continue the evaluation of EGCG as a polymicrobial biofilm therapeutic. We recommend that organic solvents, such as DMSO and dimethylformamide, would be suitable for further research [50]. In addition, the effect of ethanol itself on polymicrobial biofilms must be investigated further, with the potential for implications in ineffectiveness of alcohol wipes, hand sanitisers, and other disinfectants.

Lastly, to continue to expand this field, an examination of confirmed and novel antimicrobial compounds in combination with differing antibiotics, as well as our cellulase treatment, must be conducted. The lipid-soluble derivative of EGCG, epigallocatechin-3-gallate stearate (EGCG-S), is a confirmed antimicrobial compound that has been demonstrated to have greater stability and lipid membrane binding capabilities than EGCG [80]. However, it has yet to be tested in polymicrobial biofilms or in combination with non-antibiotic molecules, serving as a suitable compound for further investigation. Additional combinatorial antimicrobials for study include DNase, proteases, bacteriophages, quorum sensing inhibitors, biofilm dispersion molecules, and flavonoid compounds [81, 82, 83, 84]. Once identified, these compounds may be utilised in combination with different classes of antibiotics to determine the most optimal treatment for biofilm eradication.

## VII. Conclusion

This study aimed to provide evidence that novel combinatorial therapies can enhance the efficacy of antibiotics in treating multispecies biofilms. Based on the crystal violet assays, CFU counts, statistical analyses, and subsequent discussions, it can be concluded that cellulase significantly reduces both biofilm biomass and viable bacterial loads in combination with ampicillin, while EGCG fails to do so.

While this provides significant evidence, the applicability of our conclusions to *in vivo* medical practices is limited by unknown species, a lack of immune factors, and potentially reduced absorbance and CFU values for cellulase treatments. Although these limitations must be considered, they are arguably diminished due to a multitude of factors.

With minor limitations, our study demonstrates the effectiveness of cellulase in enhancing antibiotic efficacy. However, there is some ambiguity regarding the ineffectiveness of EGCG, raising the question of to what extent this ineffectiveness is determined by species-specificity or the ethanol solvent.

Our research has helped to address a fundamental gap within the literature, the effectiveness of treating polymicrobial biofilms with combinatorial therapies. Due to the high prevalence within bacterial infections, including chronic wounds, and increased levels of antibiotic resistance, the need to continue to investigate combinatorial treatments of polymicrobial biofilms is of the utmost importance. We recommend that future investigations focus on gathering further evidence of the roles cellulase and EGCG play in multispecies biofilm treatments, developing novel combinatorial therapies for polymicrobial biofilms, and conducting clinical trials of successful studies to examine *in vivo* effects, resulting in the introduction of more combinatorial antibiofilm treatments to the pharmaceutical market. Carrying out this work, the catastrophic rise of antibiotic resistance will be contested, reducing the disease burden, consequently saving the lives of millions of individuals.

