## Supplemental Data Tables 1-6 for "An Investigation of Cellulase and Epigallocatechin-3-gallate as Enhancers of Antibiotic Efficacy in Polymicrobial Biofilm Treatments"

### IX. Appendix

**Table 1. Full crystal violet spectrophotometry absorbance data set.** Biological repeat averages utilised in Figure 6. Technical replicates not depicted due to identical values.

| Treatments | Biological Repeat 1 | Biological Repeat 2 | Biological Repeat 3 | Average |
|---|---|---|---|---|
| Control | 0.162 | 0.181 | 0.186 | 0.176 |
| Ampicillin | 0.194 | 0.208 | 0.2 | 0.201 |
| Cellulase | 0.079 | 0.13 | 0.141 | 0.314 |
| EGCG | 0.257 | 0.33 | 0.356 | 0.117 |
| Ethanol | 0.244 | 0.269 | 0.342 | 0.285 |
| Cellulase + EGCG | 0.153 | 0.105 | 0.117 | 0.125 |
| Cellulase + Ampicillin | 0.162 | 0.186 | 0.151 | 0.428 |
| EGCG + Ampicillin | 0.402 | 0.428 | 0.455 | 0.166 |
| Cellulase + EGCG + Ampicillin | 0.172 | 0.254 | 0.234 | 0.220 |

**Table 2. Full Tukey's Test statistical data set for crystal violet spectrophotometry.**

For information regarding the letters utilised to denote treatments, please visit **Table 3.**

| Treatments | Q statistic | *p*-value |
|---|---|---|
| A vs B | 1.264 | 0.900 |
| A vs C | 3.099 | 0.452 |
| A vs D | 7.169 | 0.002 |
| A vs E | 5.645 | 0.019 |
| A vs F | 2.667 | 0.617 |
| A vs G | 0.520 | 0.900 |
| A vs H | 13.090 | 0.001 |
| A vs I | 2.268 | 0.767 |
| B vs C | 4.364 | 0.110 |
| B vs D | 5.905 | 0.013 |
| B vs E | 4.381 | 0.108 |
| B vs F | 3.931 | 0.188 |
| B vs G | 1.784 | 0.900 |
| B vs H | 11.826 | 0.001 |
| B vs I | 1.004 | 0.900 |
| C vs D | 10.268 | 0.001 |
| C vs E | 8.744 | 0.001 |
| C vs F | 0.433 | 0.900 |

| Treatments | Q statistic | *p*-value |
|---|---|---|
| C vs G | 2.580 | 0.650 |
| C vs H | 16.190 | 0.001 |
| C vs I | 5.368 | 0.028 |
| D vs E | 1.524 | 0.900 |
| D vs F | 9.835 | 0.001 |
| D vs G | 7.688 | 0.001 |
| D vs H | 5.922 | 0.013 |
| D vs I | 4.900 | 0.054 |
| E vs F | 8.311 | 0.001 |
| E vs G | 6.164 | 0.009 |
| E vs H | 7.446 | 0.001 |
| E vs I | 3.377 | 0.350 |
| F vs G | 2.147 | 0.813 |
| F vs H | 15.757 | 0.001 |
| F vs I | 4.935 | 0.051 |
| G vs H | 13.610 | 0.001 |
| G vs I | 2.788 | 0.571 |
| H vs I | 10.822 | 0.001 |

**Table 3. Legend for Appendix Table 2 Crystal Violet Treatments.**

| Letter | Treatment |
|---|---|
| A | Control |
| B | Ampicillin |
| C | EGCG |
| D | Cellulase |
| E | Ethanol |
| F | EGCG + Cellulase |
| G | EGCG + Ampicillin |
| H | Cellulase + Ampicillin |
| I | Cellulase + EGCG+ Ampicillin |

**Table 4. Full colony forming unit counts data set.** Counts of between 20 and 200 were utilised for analysis with the exception of Cellulase + Ampicillin Biological Repeat 1 with a dilution of $10^{-5}$. Biological repeats were averaged for calculations. Technical repeats are not depicted due to identical counts. To maintain three replicates per sample, we include the 18 colony count for the cellulase + ampicillin treatment. The abbreviation "TMTC" denotes that there were "too many to count" in terms of colonies.

| Treatments | Biological Repeat | $10^{-1}$ | $10^{-2}$ | $10^{-3}$ | $10^{-4}$ | $10^{-5}$ | $10^{-6}$ | $10^{-7}$ |
|---|---|---|---|---|---|---|---|---|
| Control | 1 | TMTC | TMTC | TMTC | TMTC | 57 | 9 | 0 |
| | 2 | TMTC | TMTC | TMTC | TMTC | 68 | 7 | 2 |
| | 3 | TMTC | TMTC | TMTC | TMTC | 62 | 6 | 0 |
| Ampicillin | 1 | TMTC | TMTC | TMTC | TMTC | 53 | 4 | 0 |
| | 2 | TMTC | TMTC | TMTC | TMTC | 66 | 4 | 1 |
| | 3 | TMTC | TMTC | TMTC | 256 | 45 | 2 | 2 |
| EGCG + Ampicillin | 1 | TMTC | TMTC | TMTC | TMTC | 55 | 5 | 1 |
| | 2 | TMTC | TMTC | TMTC | TMTC | 73 | 0 | 0 |
| | 3 | TMTC | TMTC | TMTC | TMTC | 81 | 12 | 1 |
| Cellulase + Ampicillin | 1 | TMTC | TMTC | TMTC | 220 | 18 | 0 | 0 |
| | 2 | TMTC | TMTC | TMTC | 209 | 27 | 0 | 0 |
| | 3 | TMTC | TMTC | TMTC | 225 | 28 | 5 | 0 |
| Cellulase + EGCG + Ampicillin | 1 | TMTC | TMTC | TMTC | TMTC | 54 | 5 | 0 |
| | 2 | TMTC | TMTC | TMTC | 230 | 36 | 2 | 0 |
| | 3 | TMTC | TMTC | TMTC | 247 | 25 | 2 | 0 |

**Table 5. Calculated colony forming unit values derived from original data set.** Averages utilised in Figure 7.

| Treatments | Biological Repeat | CFU/mL | Average |
|---|---|---|---|
| Control | 1 | 114000000 | 124666667 |
| | 2 | 136000000 | |
| | 3 | 124000000 | |
| Ampicillin | 1 | 106000000 | 109333333 |
| | 2 | 132000000 | |
| | 3 | 90000000 | |
| EGCG + Ampicillin | 1 | 110000000 | 139333333 |
| | 2 | 146000000 | |
| | 3 | 162000000 | |
| Cellulase + Ampicillin | 1 | 36000000 | 48666666.7 |
| | 2 | 54000000 | |
| | 3 | 56000000 | |
| Cellulase + EGCG + Ampicillin | 1 | 108000000 | 76666666.7 |
| | 2 | 72000000 | |
| | 3 | 50000000 | |

**Table 6. Full Tukey's Test statistical data set for CFUs.** For information regarding the letters utilised to denote treatments, please visit **Table 6**.

| Treatment | Q-statistic | *p* value |
|---|---|---|
| Control vs Ampicillin | 1.245 | 0.895 |
| Control vs EGCG and Ampicillin | 1.195 | 0.900 |
| Control vs Cellulase and Ampicillin | 6.194 | 0.009 |
| Control vs EGCG, Cellulase, and Ampicillin | 3.912 | 0.112 |
| Ampicillin vs EGCG and Ampicillin | 2.445 | 0.461 |
| Ampicillin vs Cellulase and Ampicillin | 4.944 | 0.036 |
| Ampicillin vs EGCG, Cellulase, and Ampicillin | 2.662 | 0.385 |
| EGCG Ampicillin vs Cellulase and Ampicillin | 7.389 | 0.003 |
| EGCG and Ampicillin vs Ampicillin, EGCG, and Cellulase | 5.107 | 0.030 |
| Cellulase and Ampicillin vs Ampicillin, EGCG, and Cellulase | 2.282 | 0.520 |
